# Phylogenetically conservative trait correlation: quantification and interpretation

**DOI:** 10.1101/2023.02.05.527214

**Authors:** Mark Westoby, Luke Yates, Barbara Holland, Ben Halliwell

## Abstract

1. Correlation across species between two quantitative traits, or between a trait and a habitat property, can suggest that a trait value is effective in sustaining populations in some contexts but not others. It is widely held that such correlations should be controlled for phylogeny, via phylogenetically independent contrasts PICs or phylogenetic generalised least squares PGLS.
2. Two weaknesses of this idea are discussed. First, the phylogenetically conservative share of the correlation ought not to be excluded from consideration as potentially ecologically functional. Second, PGLS does not yield a complete or accurate breakdown of A-B covariation, because it corresponds to a generating model where B predicts variation in A but not the reverse.
3. Multi-response mixed models using phylogenetic covariance matrices can quantify conservative trait correlation CTC, a share of covariation between traits A and B that is phylogenetically conservative. Because the evidence is from correlative data, it is not possible to split CTC into causation by phylogenetic history versus causation by continuing reciprocal selection between A and B. Moreover, it is quite likely biologically that the two influences have acted in concert, through phylogenetic niche conservatism.
4. Synthesis: The CTC concept treats phylogenetic conservatism as a conjoint interpretation alongside ongoing influence of other traits. CTC can be quantified via multi-response phylogenetic mixed models.

## Introduction

Ecological research often takes an interest in correlations across species between two traits, or between a trait and a property of the species habitat. For example, seed size is correlated (fairly loosely, r^2^ = 0.29) with the size reached by species as adults (Falster, Moles, and Westoby 2008). One motivation for investigating how closely traits are correlated is simply to understand variation across the world’s species, and to quantify how traits might be clustered together into spectra of variation. For example, a unified size spectrum has been suggested (e.g. Díaz et al. 2016) that embraces both seed size and adult size. Another motivation is that an observed correlation might be consistent with some proposed mechanism connecting the two traits, or alternatively a lack of correlation might argue against a mechanism. For example, it can be suggested that taller species typically suffer more competitive mortality between seedling and reproductive stages, and this puts a stronger selective premium on large seed size (Falster, Moles, and Westoby 2008).

A present-day correlation between seed size and potential plant size across species can be interpreted as caused by trajectories of change through past evolution. Equally, the past trajectories can be interpreted as movement toward evolutionary attractors, produced by an ecological mechanism that exerts continuing selective pressure in the present day. Either of those versions of causation are consistent with observed correlations between traits, or between a trait and habitat.

It is important to be clear that correlations across species come from observational or survey evidence. They can offer support for some proposed mechanism or argue against it, but they can not significance-test them in the same sense as manipulative experimental treatments can. In manipulative experiments, the treatment is cause and the outcome is effect, and other factors are controlled or randomized so that each replicate yields an independent item of evidence for the link between cause and effect. Because the items of evidence are independent, a P-value for the ensemble of events can be calculated with confidence. Whereas in survey evidence, some unmeasured or uncontrolled variable might be creating a correlation between the two focal traits, or counteracting it.

Where investigators have thought about third and fourth variables as possible influences, and have been able to obtain measurements for them, a more limited sort of independence can be obtained by controlling or partialling for these third or fourth variables, or equivalently by applying multiple regression. Residuals are obtained for the focal variables after regression on the covariates, and correlations between the residuals are then inspected. But this is still a very different sort of independence compared to the evidence that emerges from a manipulative experiment. If an A-B correlation disappears after partialling for C, it still remains a possibility that C was a secondary correlate and the true mechanism runs between A and B. Plus there remain variables D, E, F and so forth that might have been the true cause but were not measured or not even thought of.

Structured causal modeling SCM (Pearl 2009) or graphical causal modeling (Cronin and Schoolmaster Jr. 2018) is a framework that purports to determine cause-and-effect relationships from observational data (Arif and MacNeil 2022). However, the conditions for identifying causation unambiguously are stringent. The causal maps are required to be directed acyclic graphs (DAGs), with no recursion to variables earlier in the causal chain. It must be possible to list all competing causal hypotheses in order to compare them, and each must correspond to a different chain of causation between variables. In our opinion (contra Cronin and Schoolmaster Jr. 2018), these conditions are not ever met by the situations of interest here, coordination across species among traits and habitat and their relationship to phylogeny. Coordination between traits happens because the current value of each trait influences natural selection on the other (recursion). Traits also influence the habitats where the clade is successful, and habitat in turn exerts natural selection on the traits (again recursion). A map leading from clade membership to trait values always has alternative causal interpretations: (1) that traits are intrinsically slow to change so that clade signal remains, and (2) niche conservatism, that there is continuing ecological selection from other traits in combination with habitat.

It is widely held that correcting or controlling or accounting for phylogeny (methodology summarized in Box 1) should be mandatory when ecologists consider present-day functionality of traits in combination with each other or with different environments (e.g. Losos 2011; Garamszegi 2014; Swenson 2020; Revell and Harmon 2022). Reviewers and editors of ecology journals commonly require authors to control for phylogeny. Despite this strong majority view insisting on the practice during review, experts have raised substantial questions about what is achieved by controlling for phylogeny (Box 2).

Correcting an A-B relationship for phylogeny uses the same logic as partialling it for a continuous variable C. The commonest justification why phylogenetic correction should be mandatory is to say that related species are not independent (Felsenstein 1985, and very often repeated up to the present day, e.g. Symonds and Blomberg 2014). An A-B relationship controlled for phylogeny is often interpreted as a corrected or improved version of the simple cross-species relationship. This interpretation is not correct. Rather, phylogenetically controlled relationships measure different properties of the data, compared to relationships across present-day species. They address a different question (see section below “What does phylogenetic generalised least squares quantify?”).

A statistical method corresponds to a generating model. Its equations, variables and probability distributions express models for causation or for prediction. Only if the generating model is well aligned with a biological hypothesis will a clear answer be delivered. The statistical models fitted, the causal or predictive maps hypothesized, and the biological questions of interest are all aspects of the same issue.

The PGLS and PIC methods mainly used for controlling for phylogeny correspond to particular generating models. Our main aim here is to show that these generating models do not necessarily correspond to questions that ecologists want to ask. Further, the fact that they are couched in terms of least squares regression of A against B does not adequately represent a generating process whereby A and B reciprocally influence each other. Another aim here is to put forward multi-response or multivariate phylogenetic mixed models (MR-PMM). These treat A and B as joint responses and partition the different correlations in a way that does not treat phylogeny and present-day function as alternative interpretations. MR-PMM are not new (Lynch 1991; Housworth, Martins, and Lynch 2004), but have not come into common use in ecology.

### Quantifying conservative trait correlation via multi-trait response models

The most straightforward reason why controlling for phylogeny should not be interpreted as automatically correcting or improving an A-B relationship, is that present-day influence from B and phylogenetic conservatism overlap as explanations for variation in A. Controlling for phylogeny is advocated with a view to discarding, or partialling out, A-B covariation that is phylogenetically conservative from the A-B relationship. From the perspective of understanding present-day ecological differences across species, this means that differences between major clades are downweighted as contributors. To the extent differences between major clades are important in present-day ecology, it risks throwing the baby (or large parts of it) out with the bathwater (Hansen 2014; de Bello et al. 2015).

A constructive solution to this problem of interpretation lies in multi-response phylogenetic mixed models (MR-PMM; Halliwell, Yates, and Holland 2022). These models decompose trait-level covariance and variance into phylogenetic and independent components (details in Box 3 and Table 1). A component of A-B correlation that is also phylogenetically structured can be identified and quantified. We refer to this quantity as the conservative trait correlation CTC. In these MR-PMM, as in PGLS, a matrix of covariances expected from a phylogenetic generating model appears as part of the residual structure on the right hand side. The key difference from PGLS is that traits A and B are jointly modelled as response variables on the left hand side (hence the name multi-response), and both their phylogenetic and independent correlations are parameters to be estimated. This makes it possible to decompose the A-B correlation into a component that is also phylogenetically structured (conservative trait correlation) and a component that is independent of phylogeny (Table 1). It has also the effect of treating the A-B relationship as a question of how they are coordinated rather than as a question of how B predicts A, analogous to standardized major axis (SMA) relationships rather than to ordinary least squares (OLS) regression (Warton et al. 2006). This will be appropriate for most evolutionary questions, since selective influences between traits or between a trait and a habitat property will be reciprocal.

**Table 1.** Where variation is attributed by the multi-response phylogenetic mixed model (MR-PMM) described here. Key parameters estimated are four standard deviations 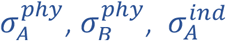, and 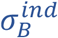, and two correlations 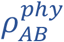 and 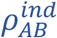.

| Components of variation | expression |
| --- | --- |
| total variation in A | $\sigma_A^2 = (\sigma_A^{phy})^2 + (\sigma_A^{ind})^2$ |
| phylogenetic signal in A | $\lambda_A = \frac{(\sigma_A^{phy})^2}{\sigma_A^2}$ |
| total variation in B | $\sigma_B^2 = (\sigma_B^{phy})^2 + (\sigma_B^{ind})^2$ |
| phylogenetic signal in B | $\lambda_B = \frac{(\sigma_B^{phy})^2}{\sigma_B^2}$ |
| A-B correlation associated with phylogeny (called here conservative trait correlation CTC) | $\rho_{AB}^{phy} = \frac{\Sigma_{AB}^{phy}}{\sqrt{\Sigma_{AA}^{phy} \Sigma_{BB}^{phy}}}$ |
| A-B correlation independent from phylogeny | $\rho_{AB}^{ind} = \frac{\Sigma_{AB}^{ind}}{\sqrt{\Sigma_{AA}^{ind} \Sigma_{BB}^{ind}}}$ |
| total A-B correlation<br>(expression in terms of lambda as given by Housworth, Martins, and Lynch 2004) | $\rho_{AB} = \frac{\Sigma_{AB}^{phy} + \Sigma_{AB}^{ind}}{\sigma_A \sigma_B} = \rho_{AB}^{phy} \sqrt{\lambda_A \lambda_B} + \rho_{AB}^{ind} \sqrt{(1 - \lambda_A)(1 - \lambda_B)}$ |

From a statistical point of view, conservative trait correlation CTC is A-B covariation where for each trait, phylogeny and the other trait jointly are associated. It is not possible to separate them. From the point of view of interpreting biological mechanism, it is quite likely that phylogeny and each trait have acted in concert on the other trait, via phylogenetic niche conservatism (next section). MR-PMM identifies phylogenetically-conservative covariation between traits A and B (Table 1), and remains agnostic whether this covariation should be attributed to phylogenetic history or to continuing reciprocal selection between the traits, or to the synergy between those two, known as niche conservatism. This is more constructive than the PGLS partitioning, which is used with a view to separating phylogenetically- conservative covariation from the estimate of the A-B relationship (see also section on PGLS).

### Phylogenetic niche conservatism

Should the present-day pattern of trait-combinations across species be interpreted as caused by trajectories of change through past evolution? Or should the past trajectories be interpreted as movement toward evolutionary attractors, which continue to be attractors in the present day? A correlation between traits, or between a trait and habitat, can be interpreted in either of these ways. Traits of ecological importance are expected often to evolve in a phylogenetically conservative way. If a new ecological opportunity or niche arises, successful occupants are most likely to be drawn from clades that already possess appropriate trait-combinations. Descendants from a clade are most likely to be successful in habitats or ways of life similar to those the clade is already adapted for. Through this phylogenetic niche conservatism, large shares of present-day adaptation and phylogeny can often be bound together as a unified causal process. Differences between major clades are often important contributors to the observed variation across ecological strategies. Phylogenetic history and present-day ecological competence are complementary explanations, not mutually exclusive alternatives.

Consider the simulations described in Fig 1. In Fig 1a there is an overall correlation between traits A and B, but the correlation is generated from the difference between two major clades, and no correlation has been simulated within each clade. A similar pattern was shown in Felsenstein’s (1985) Fig 7. His interpretation was that “It can immediately be seen that the apparent relationship ….. is illusory”. However, Price (1997) showed that a similar pattern could in fact be produced by continuing selective forces. Fig 1b illustrates this, using a simulation driven by the same principles as Price.

**Figure 1.**
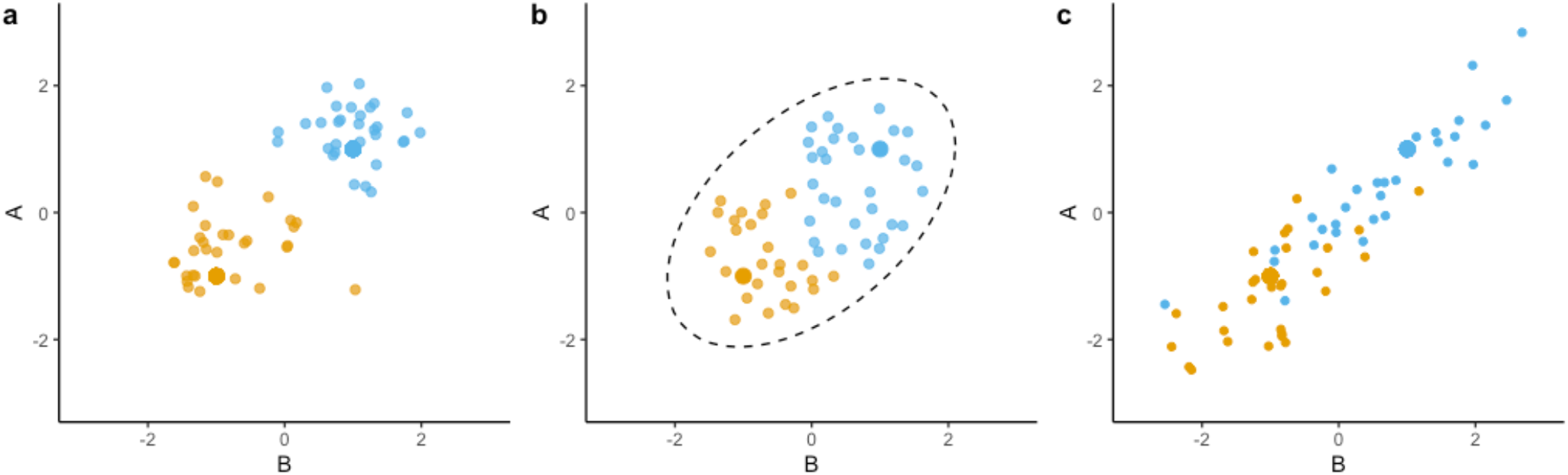
Data simulated under different evolutionary models, beginning from two clades (orange and blue) separated in a space described by two traits A and B. Large solid points represent the most recent common ancestor for the orange and blue clades in each simulation. Species then radiate, and traits diversify, within each clade. In simulation (a), radiation of each major clade proceeds by Brownian motion. The overall correlation between traits has been produced entirely by the starting points of the two major clades. In simulation (b), the radiations are positioned at random within a region of trait space (broken line) whereby only trait combinations within the line are competent to support viable populations. New viable species are more likely to arise from clades that have existing species nearer to them in trait space. The observations in present-day species are not distinguishable between simulations (a) and (b), illustrating how historical vs present-day determination of an overall correlation can often not be distinguished by analysis of present-day data. A version of this comparison was first given by Price (1997). In simulation (c), data are produced from a MR-PMM. Positive correlations between A and B are operating on both the phylogenetic and independent level 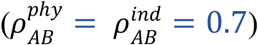, resulting in a tight overall relationship between A and B. Importantly, correlations within each of the major clades and also between them, are both important drivers of the present-day spectrum of variation.

In Fig 1b the broken line circumscribes an ecological attractor, a region of trait combinations in niche space that are ecologically competent. The shape of this region is of high interest for ecologists. Indeed, this is the motivation for looking at scattergrams of one trait vs another. It is supposed for both (a) and (b) of Fig 1 that orange and blue symbols represent sister clades that have diverged in trait space toward lower left and upper right. Panel (a) then assumes Brownian motion, while (b) assumes that new species can emerge only within the viable trait- space and tend to be drawn from the existing clade that is nearest in trait space. In both cases each clade is phylogenetically conservative. In (a) the conservatism takes the form of sluggish Brownian motion. (If the Brownian motion is rapid, then the historical difference between the two major clades is quickly washed out.) In (b) conservatism arises from a constrained range of ecological possibilities.

The point is that the observed pattern across present-day species cannot help to decide which of these causative interpretations is more likely. Further, the process in Fig 1b is both phylogenetically conservative and also caused by ecological constraints continuing into the present day. Data analysis should not treat these as competing alternatives. Better for it to identify conservative trait correlation, the share of trait correlation that might be attributed either to phylogenetic history or to continuing functionality or to a combination of the two.

The question how much to interpret functional traits in terms of past history versus in terms of present-day competence itself has a history (brief summary in Box 4). To some extent it reflects tension between the outlooks of evolutionists and ecologists.

When ecological selection has favoured high trait A in conjunction with high trait B through the length of phylogenetic history, as well as in the present day (as in Fig 1c), then ordinary regression across species and PGLS will yield similar results (Fig 2, simulation S1), because the trait correlation pattern across phylogenetic divergences is similar to the pattern across present-day species. This is a very common case in real datasets (Ackerly 1999; Carvalho, Diniz-Filho, and Bini 2006). Nevertheless, this similarity should not be the basis for choosing OLS in preference to PGLS or vice versa. These two analyses, and also MR-PMM, are different in what features of the data they model. Analyses should be chosen to match the assumptions of the generating model and the question being addressed, even though OLS and PGLS quite often yield similar slopes.

**Figure 2.**
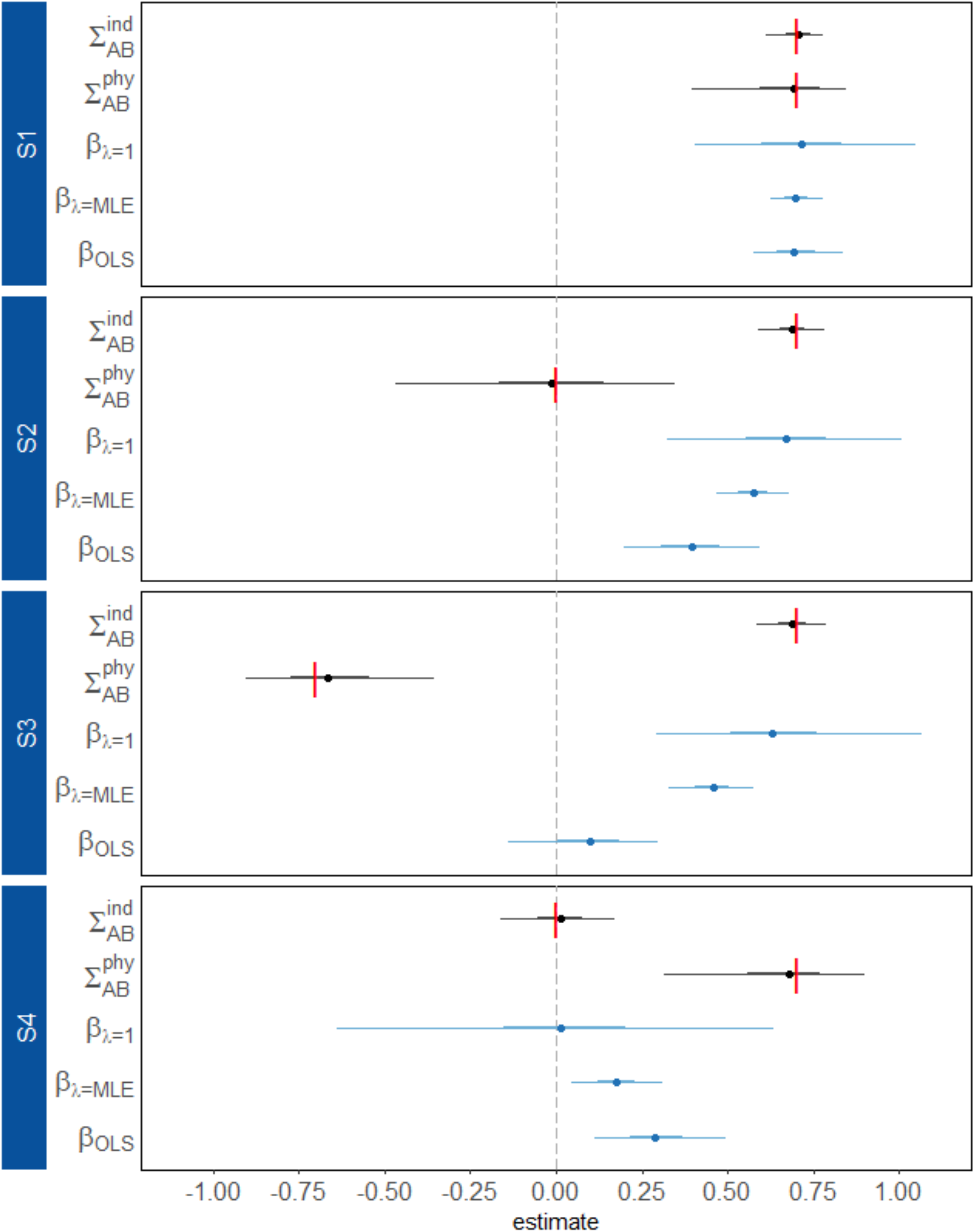
Parameter estimates for phylogenetic and independent covariances (black) from MR-PMM, and beta coefficients (blue) from PGLS (β_*λ*=1_), PGLS with lambda optimised (β_*λ*=MLE_), and OLS (β_OLS_) fit to simulated datasets (S1-4). Points represent the median posterior estimate across 400 model fits, with heavy and light wicks showing the 50% and 90% sample quantiles, respectively. For each simulation, 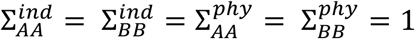. True values for 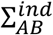 and 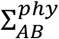 used to generate the data for each simulation are indicated by vertical red bars.

### Controlling for phylogeny does not confer strong-sense independence

The most common justification why controlling for phylogeny should be mandatory is to say that related species are not independent (Felsenstein 1985, and very often repeated up to the present day). This justification implies that independence is restored by controlling for phylogeny. But that implication is only correct in a very limited sense.

Independence is used with two meanings. The looser meaning is simply “uncorrelated”. Seed size can be said to lack independence from plant asymptotic height. Independence can be said to be restored by replacing the absolute seed size with residuals around a regression of seed size on plant height. However, this is a conditional independence, from a fitted function of plant height only, not from all possible confounding variables.

The tighter meaning of independence is about formally significance-testing a hypothesized causal mechanism. If causal events are independent, their probabilities can be multiplied to yield the probability of the ensemble of events. When phylogenetic correction is said to be obligatory because cross-species correlations lack independence, the suggestion is that after correction, independence will be ensured in this tighter sense, and a reliable significance test will ensue. But significance tests from survey or correlative data should not be interpreted as clean tests of causation anyhow. An A-B correlation may be more or less consistent with a proposed causation, but it does not provide significance-tested proof. Correcting for phylogeny using structured residuals addresses one sort of cross-correlation, but not all.

There may always be further variables unmeasured or not even thought of that are correlated with both A and B (Price 1997). And further, in a network of interconnected variables, correlations do not by themselves resolve the pathways along which causation runs.

Criteria need to be invoked from outside the correlative data. Some causal pathways might seem more plausible than others in light of known mechanisms, or parsimony can be invoked in choosing among statistical models, or combinations of plausibility with parsimony (Yates, Richards, and Brook 2021).

Independence in this sense of independent events showing causality can really only be assured in manipulative experiments. Treatments come before outcomes in an experiment’s timeline, so it is clear which is cause and which is effect. And factors other than treatments are randomized or physically controlled, so each replicate becomes definitively an independent instance of a treatment giving rise or failing to give rise to an outcome. In situations such as social science surveys or comparisons across present-day species, there is no way to assure independence in this rigorous sense (Hernán, Hsu, and Healy 2019).

Significance tests should not be taken too literally when analysing survey data, but r^2^ and similar indices that quantify the strength of correlations are useful descriptors. Cross-species relationships are correlations not causation, and remain so after adjusting for phylogeny.

To invoke independence in the context of estimating a P-value you need to specify what process is being tested for. Then the question is whether two or more events or links between variables are independent <u>as evidence for that process</u>. Independence is a property of the hypothesis as well as of the data structure. To say that past divergences are independent cases for a link between traits A and B while present-day species are not, is just another way of asserting that past divergences are a legitimate causative explanation while continuing present-day selection is not (Box 4).

Arguments over the primacy of causal processes cannot be resolved from data that are cross-correlated. We should look to statistical methodologies that offer the most informative decomposition of trait variance, without preferencing one causality over another. Causality can then be interpreted in light of knowledge about physiological mechanisms, or manipulative experiments demonstrating how particular trait values confer advantage depending on other traits or on habitat.

### What does phylogenetic generalised least squares quantify?

Saying that PGLS accounts for phylogeny does not tell us what it actually is. What is measured by the PGLS slope and confidence intervals, and how should it be interpreted?

For phylogenetically independent contrasts PICs and for PGLS with λ set at 1, the slope coefficient with associated confidence interval describes divergences in A as predicted from divergences in B, taken across the population of all past divergences inferred at all the nodes in the tree. The significance test (whether the confidence intervals on the slope span zero) assesses consistency, the question whether trait divergences were correlated across most or all of the nodes.

The interpretation of PGLS with λ estimated lies somewhere between the extreme cases of λ = 0 (OLS) and λ =1 (original PGLS). Fitted λ with a value less than 1 but still not zero can be interpreted as a rescaling of branch lengths in the phylogeny (Symonds and Blomberg 2014). Low lambda elongates the terminal branches, reducing the expected similarity between related species. With fitted λ the PGLS slope likely still reflects an ensemble of slopes across all divergences, but with divergences calculated on a tree with branch lengths modified by λ. PGLS assumes that the true generating process is consistent with ‘B predicts A’ and that B has no phylogenetic signal. These assumptions are usually not true. Unless predictor B actually is free of phylogenetic signal, the slope is confounded with the phylogenetic component of the residual variance (Supplementary 2, see also simulations in next section). This confounding will commonly have the result that some but not all of the phylogenetic signal in the A-B relationship remains in the slope estimate. Analogous issues occur in spatial statistics where environmental predictors with spatial signal are confounded with the spatial component of the residual variance (Marques, Kneib, and Klein 2022; Warton 2022).

The focus of PICs and PGLS on describing correlation in past divergences accords with the insistence of evolutionists that adaptation is defined as the selective circumstances when a trait or trait value first came about (Box 4). PGLS does not directly measure the relationship between traits in the present day, unless λ = 0 when it becomes an ordinary least squares regression. It is the nature of most trees that there are many nodes near the tips and rather few deep in the tree. As a result, deep nodes have only a minor influence on the PGLS-estimated relationship between divergence in A and divergence in B. But the consequences of a single deep divergence for the pattern across present-day species can sometimes be very substantial (Fig 1). Consider for example the divergence between angiosperms and gymnosperms. As well as qualitative differences such as tracheids vs vessels for water transport, these two major clades of seed plants have widely different strategies with regard to quantitative traits such as vein density in the leaves, seed size and leaf mass per area (e.g. Ackerly and Reich 1999; Brodribb et al. 2005; Díaz et al. 2016). This only counts as one divergence among many in a PGLS, but it has large consequences in terms of ecological strategies sustained in the present day.

The relationships quantified by OLS and by PGLS are different, OLS a pattern across present-day species, PGLS a pattern across past evolutionary divergences (at least with λ = 1). Biologically, these are naturally complementary questions, but they are different, and one should not be seen as replacing the other. MR-PMM quantifies both types of relationship within a single analysis, but not in quite the same way, since it models phylogenetic signal in both traits and formulates the relationship as A-B coordination rather than as predicting A from B.

### Simulations to illustrate how MR-PMM compares to PGLS

To illustrate issues around niche conservatism and the decomposition provided by MR-PMM, we simulated datasets of two traits A and B. For each simulation, A and B were given the same independent or residual covariance, but different phylogenetic covariances (Supplementary 1 for details). We simulated 400 replicate datasets by generating random pure birth trees of 200 taxa. The two traits A and B were simulated from the full cross- covariance structure of the MR-PMM, comprising phylogeny-independent covariances crossed with phylogenetic covariances, rather than generating A as a scalar multiple of B, as is assumed in PGLS (e.g. Revell 2010). This structure allows the independent and phylogenetic variance in A and B to be defined separately and explicitly.

MR-PMM successfully recovers the trait-level covariances of each generating model (Fig 2), as expected since it corresponds to the generating model. Comparing these covariance estimates with PGLS is more complicated since PGLS is a single-response model and does not report covariances directly, only the β slope coefficient. To facilitate comparisons, the independent and phylogenetic variances of each trait were set to one which places the slope coefficients on the same scale as the correlation coefficients (see Supplementary 1 for details). This choice of scale means that the β_OLS_ estimate is approximately equal to the mean of the simulation values for independent and phylogenetic correlation components, β_*λ*=1_ is approximately equal to independent component, and β_*λ*=MLE_ attains an intermediate value depending on the estimated λ.

These simulations illustrate the following. First, when covariances between traits are similar on the phylogenetic and independent level (Fig 2, S1), then β_*λ*=1_, β_*λ*=MLE_ and β_OLS_ are also similar. Biologically, this is a common outcome (Price 1997; Ackerly 1999; Carvalho, Diniz- Filho, and Bini 2006). Second, β_*λ*=1_ has the effect of disregarding covariance associated with phylogenetic history during calculation of the β coefficient. Our central point in this paper is that phylogenetically-associated covariance should not be automatically set aside, because niche conservatism is both phylogenetic and also represents selective attractors that continue into the present day (Figure 1). Third, in the extreme case where the correlation between a pair of traits occurs exclusively on the phylogenetic level, β_*λ*=1_ is likely to report no relationship (Figure 2, S4). To the extent that differences between major clades are important in present-day ecology, this result represents a false negative. Fourth, optimising *λ* does not resolve this problem, rather it represents a compromise between the assumptions of PGLS and OLS. Finally, because β_*λ*=1_, β_*λ*=MLE_ and β_OLS_ are all products of single-response models, they represent single-number summaries of the different components of A-B covariance present in the data (Supplementary Information 2). This means they are poor approximations of the true generating model when phylogenetic and residual covariances differ in sign (S3) or even magnitude (S2).

## Conclusion

Both evolutionary and ecological questions about traits are important, but they are not the same. For ecologists interested in the present-day relationship of traits to habitat or each other, phylogenetic correction has been justified largely from the perspective that trait correlation across species might be misleading. This formulation is missing the point from the outset. Correlative data are undoubtedly capable of being misleading, and need to be approached with that mindset. But it is wrong to think that controlling or accounting for phylogeny obviates the problem.

Phylogenetically independent contrasts PICs ask about the history of divergences at nodes. The divergences, rather than the present-day species, are the population of interest. The question whether divergences in A have been consistently associated with divergences in B is a natural one for evolutionists to ask. It is complementary to the ecological question about trait-combinations that are competent in the present day, but it is not the same question.

Phylogenetic generalized least squares PGLS is currently widely recommended and used. When used with *λ*= 1, it is mathematically equivalent to PICs. However, the actual historical divergences are not inspected or graphed as they are for PICs. The slope estimate with *λ*= 1 describes the power of divergence in B to predict divergence in A, across the ensemble of divergences or nodes. As was the case for PICs, this slope estimate answers an interesting question, but not the question how traits are related across present-day species.

When PGLS is used with *λ* fitted to the data, *λ* will usually lie intermediate between 0 and 1, since for most traits there is some phylogenetic signal but not perfect correlation with a phylogenetic generating model. The strength of the residual phylogenetic influence is then measured via *λ*. The estimated slope is intermediate between the slope across divergences and the slope across present-day species.

Multi-response phylogenetic mixed models open a path to interpreting covariance structure better in two ways, we believe. First, their generating model deals in A-B covariation, which reflects the nature of reciprocal influences between traits and habitat more satisfactorily than regression-style models predicting A from B. Second, they quantify the variance and covariance components more comprehensively. In particular, they quantify conservative trait correlation CTC, and remain agnostic about whether it is caused by history, by continuing evolutionary attractors, or by both. Historical and present-day accounts of causation are, in fact, complementary. Over evolutionary time, new ecological opportunities will very often have been taken up by speciation from clades that already possess a configuration of traits close to what will be most successful.

## Acknowledgments

Ian Wright kindly put the authors in touch with each other. Warm thanks also to Daniel Falster and David Warton for much discussion of this topic over the years. Halliwell and Yates were partly funded by The Australian Research Council Centre of Excellence for Plant Success in Nature and Agriculture (CE200100015).

## Boxes and Tables

### Box 1: Phylogenetic correction in brief

Consider a dataframe giving traits or habitat properties (columns) across a number of present-day species or other entities (rows). Also, the species in the data table are connected by a tree structure representing their phylogeny, as best it is known. Phylogenetic correction of correlations between columns in such a dataframe has two elements. There is a statistical procedure, and then an interpretive step whereby the phylogenetically -adjusted relationship between two traits or between a trait and a habitat is seen as corrected, compared to the raw correlations across present day species. The implication is that the phylogenetically adjusted relationship is more reliable, or more enlightening, or that the model is more complete. Statistical method and interpretation are linked. What generating process is being assumed by the statistical model, and hence what question exactly does a given statistical method ask?

One version of the statistical procedure is to transform a set of present-day species into a set of evolutionary divergences or phylogenetically independent contrasts PICs (Felsenstein 1985). At each node in the tree, an evolutionary divergence or PIC is inferred for each trait. These divergences, rather than present-day species, then become the objects under study, and the cases or items of evidence in a statistical procedure. The question is whether divergences in trait A tend to be correlated in size and direction with divergences in trait B. (For a polytomy, there is a regression between trait A and trait B across the set of descendant species or nodes. Indeed for a dichotomy, the divergences can also be thought of as a two-point regression.) The effect has something in common with a pairing design in social science, where individuals are matched for (say) gender or age or income, then differences (“contrasts”) are calculated across the pair for other variables, and the analysis proceeds using those contrasts as items of evidence, rather than the individuals themselves.

Currently the method most often used is phylogenetic generalised least squares PGLS (Grafen 1989; Martins and Hansen 1997). This is a regression model for relationships between traits across species. The expected residual covariances between each pair of species are modelled in such a way that higher covariance is expected when the species have diverged more recently on the phylogenetic tree. If two species are outliers in the same direction and also have a relatively recent common ancestor, then some covariance between them is seen as expected, and the influence of those residuals on the position of the fitted line is downweighted accordingly. In other words, the idea that traits are for unspecified reasons slow to change through evolutionary time (phylogenetic inertia) is part of the causation being modeled.

The phylogenetically expected covariances in PGLS scale with the combined branch lengths shared between species, reflecting a Brownian-motion or diffusion model for trait change. Often a parameter *λ* (Pagel 1999) is fitted by maximum likelihood as part of the model. This is a multiplication factor in the range 0 to 1 for the off-diagonal elements of the phylogenetically expected residual covariance matrix. If *λ* is near zero, this effectively makes the terminal branches of the tree very long, with little covariance expected even between sister species. (In phylogenetic mixed models PMM discussed in Box 3, an equivalent scaling is estimated for each response trait (Halliwell et al. 2022)). In addition to the basic Brownian- motion model, a variety of more complex models have been developed (overview in Garamszegi 2014), that fit parameters for rates of trait change that vary through time or in response to other variables.

PGLS with *λ* fixed to 1 is mathematically equivalent to PICs, which iteratively calculate divergences or contrasts at each node through the phylogenetic tree and treat those as a population of events (Blomberg et al. 2012; Symonds and Blomberg 2014). Under these circumstances the regression slope and confidence intervals reported by PGLS are summarizing the population of regression slopes across all the divergences or nodes in the phylogenetic tree. PGLS with *λ* = 0 yields the ordinary least squares regression slope across present-day species. With intermediate *λ*, the slope will lie somewhere in between those two meanings. Mathematical treatment is provided as Supplementary 2.

### Box 2. Selected quotes that illustrate uncertainty among experts about what is achieved when controlling for phylogeny

The majority or standard view is expressed by Garamszegi (2014) in the preface to an edited book: *“Statistically, the effect of phylogeny can be regarded as a confounding factor that violates assumptions about non-independence of the unit of analysis, and that potentially introduces spurious correlations across traits*.*”* Similarly Huey et al. (2019): *“Independent contrasts enabled comparative biologists to avoid the statistical dilemma of nonindependence of species values, arising from shared ancestry … Felsenstein (1985) rapidly and radically changed both evolutionary and organismal biology … No one would consider ignoring phylogeny when analyzing data involving multiple species …”*

As against that majority view, the following quotes make the point that adaptation to niche and phylogenetic history should not be treated as competing alternatives. Housworth et al (2004) wrote “*the heritable component contains not only genetic changes but also nongenetic contributions to the phenotype, such as environmental or cultural contributions, that are described by the phylogenetic relationship among the taxa*.”. Hansen (2014) wrote *“if related species tend to occur in similar environments (i*.*e*., *having similar values of their predictor variables), then we still expect a phylogenetic signal in the response variable. Correcting for phylogeny in this situation is throwing the baby out with the bathwater … [perhaps] the application of phylogenetic comparative methods has done more harm than good in the study of adaptation*.” De Bello et al (2015) wrote *“Phylogenetic relatedness between species should not be considered a bias to be corrected, but rather an evolutionary signal that allows results to be interpreted at different evolutionary scales*.*”*

Any given model reflects a hypothesis about processes generating the observed data (Uyeda, Zenil-Ferguson, and Pennell 2018): *“[phylogenetic comparative models] PCMs are powerful tools for drawing inferences from interspecific data but they necessarily imply some types of causal structures and negate others. It is too much to ask of our methods to decide what questions we ought to ask*.*”*

And causation can not be decisively inferred from survey data: *“the validity of causal inferences depends on structural knowledge, which is usually incomplete, to supplement the information in the data. As a consequence, no algorithm can quantify the accuracy of causal inferences from observational data”* (Hernán, Hsu, and Healy 2019).

### Box 3: Multi-response phylogenetic mixed models MR-PMM as applied to dissecting covariance between two traits across species

In multi-response phylogenetic mixed models, two or more traits appear as responses on the left hand side of the model equation. Terms on the right hand side include a matrix of covariances expected from a model of trait change through the phylogeny, as well as trait-level intercepts and possibly fixed or random variables. With respect to a single-response model, changing the status of trait B from a predictor for A to a joint response variable with phylogenetically structured residuals allows phylogenetically conservative A-B correlation (conservative trait correlation CTC) to be quantitatively identified. The multi-response approach treats the A-B relationship as a question of trait coordination rather than a question of predicting A from B. For allometric relationships, this joint view yields a consistent estimate of trait coordination, via a decomposition of their residual covariation, unlike the predictive view where the slope estimates depend on whether A is predicted from B or vice versa (Warton et al. 2006). Indeed, for data generated from a MR-PMM, slope estimates for B from a misspecified single-response model such as PGLS confound various components of the generating model (explained further in Supplementary 2).

MR-PMMs offer a sufficiently complex and more biologically appropriate model structure than their single-response analogues. These models simultaneously account for phylogenetic signal in all included traits and permit a decomposition of the estimated trait correlation according to dependence on phylogeny. For two species traits A and B, a multi-response mixed model with phylogenetic covariances modelled as a random effect takes the form

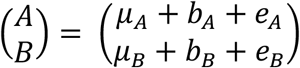

The μ’s are vectors of fixed effects, which can be any linear predictive equation. When the interest is only in the relationship between traits A and B, i.e. there are no predictors in the model, these fixed effects would contain only an intercept for each trait.

The phylogenetic random effects b_A_, b_B_ and the phylogeny-independent effects e_A_, e_B_ are drawn from multivariate normal distributions

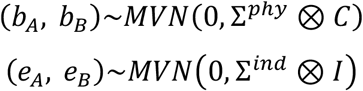

For two response traits, A and B, and n species in the phylogeny, the covariance matrices for the random effects and independent errors are of dimensions 2n x 2n. The covariance of the phylogenetic random effects ∑^*phy*^ ⊗ *C* is the Kronecker product of a 2 × 2 trait-level correlation matrix, ∑^*phy*^, with *C*, the n x n matrix of expected error covariances given a model of trait evolution applied to a phylogenetic tree. For the simplest case of Brownian motion, *C* is the phylogenetic relatedness matrix. The covariance structure of the residuals or phylogeny-independent elements ∑^*ind*^ ⊗ *I*, is the Kronecker product of a 2 × 2 trait-level correlation matrix ∑^*ind*^, with I, an n x n identity matrix (1 for diagonal elements and 0 for off-diagonal elements). For the two-trait PMM, we estimate two phylogenetic variances for A and B (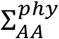 and 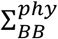) and the phylogenetic covariance between A and B 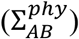. The same is true for independent (co)variances in the elements of ∑^*ind*^. When scaled by the relevant variance components, these covariances yield phylogenetic and residual correlations. Thus, when appropriately parameterized, the MR-PMM estimates each element listed in Table 1.

MR-PMM looks at the correlation between A and B rather than at predicting one from the other. Given a fitted MR-PMM, further derived quantities such as (standardized) major axes relating A to B (Warton et al. 2006) can be constructed from either point estimates or posterior distributions of the variance and covariance parameters associated with the two traits.

In principle, models with this layout can have any number of species traits or habitat properties on the left hand side, and also other predictors included in the fixed-effect terms on the right hand side. More complex models require more replication to yield reliable estimates (Housworth, Martins, and Lynch 2004). For simplicity, we have confined this explanation to the correlation between two Gaussian traits, but response variables are not required to be Gaussian distributed. See Halliwell, Yates and Holland (2022) for details including worked examples in two popular R packages, ‘MCMCglmm’ and ‘brms’.

### Box 4: Past and present-day causation

The question whether adaptation should be interpreted as a past versus a present-day process has long been debated. Palaeobiologists and evolutionists have insisted that adaptation should refer only to the selective circumstance that initially gave rise to a trait. For example, Gould and Vrba (1982) coined “exaptation” for functionality that came about subsequent to a trait’s origin, in order to reserve adaptation for functionality at the time of origin. (For a quantitative trait such as adult body size, they must have meant “origin” to refer to the time the trait arrived at a particular value.)

This defining of terms by evolutionists has mostly stuck over the ensuing 40 years. For example, Paradis (2014) wrote: *“we can define the phylogenetic comparative method as the analytical study of species, populations, and individuals <u>in a historical framework</u> with the aim to elucidate the mechanisms at the origin of the diversity of life*.*”* Losos’s (2011) presidential address to American Society of Naturalists discussed traits and phylogenies. Summarizing the history of ideas, he wrote “*the key turning point was the publication of Felsenstein’s (1985) article in the American Naturalist, which presented the issue of shared ancestry as a difficulty in comparative analysis and the independent contrasts method as the solution …. publication of books by Brooks and McLennan (1991) and Harvey and Pagel (1991) completed the revolution. Since that time, there has been a continuous, unabated rise in the development and use of phylogenetic comparative methods. Comparative studies now are essentially unpublishable unless analyzed in a phylogenetic context …*”. Losos 2011 also wrote in a footnote: “*many reviewers … have been concerned that this article will give license to ecologists and other ne’er-do-wells to ignore phylogenetic approaches entirely. So, just to be clear, I will say it again: phylogenetics is an important approach for studying historical events …. This article should not be read as license to ignore phylogenetic information in comparative studies!*”

Phylogenies are indeed essential for studying the history of divergences. But what has happened here is that comparative studies have been defined as being about history, in the same way as adaptation earlier was defined as being about history. On the other hand, ecologists have a continuing interest in the question what traits or trait-combinations make species successful in what situations in the present day. Losos intended, no doubt, to express collegiality from evolutionists toward ecologists when he breezily called them ne’er-do- wells. But the collegiality did not extend to permitting ecologists to consider adaptation and comparative studies as questions about the present day.

The essential point for ecologists is that patterns such as in Fig 1 where a trait is correlated both with another trait and with phylogenetic history, called here conservative trait correlation CTC, can potentially arise from a deep historical divergence followed by limited subsequent change, or from continuing selection in the present day. The observed pattern does not give a basis for preferring one explanation to the other, and moreover the two need not be mutually exclusive. For ecologists aiming to describe trait combinations that confer present-day competence, it is not sensible to remove the conservative trait correlation from consideration. That is why we recommend instead the partitioning of variation provided by MR-PMM (Table 1).

Supplementary 1: Simulations

Supplementary 2: Mathematical aspects of phylogenetic mixed models

Supplementary 3: R code for the simulations reported in Figs 1 and 2

