## Supplementary 3 for "Phylogenetically conservative trait correlation: quantification and interpretation"

#-------------------------------------------------------------------------------------#
#                                                                                     #
# Phylogenetically conservative trait correlation: quantification and interpretation  #
#                                                                                     #
# Supplementary Material 3                                                             #
#                                                                                     #
#-------------------------------------------------------------------------------------#


library(geiger);library(ape);library(phylolm);library(phangorn);library(phytools)
library(brms);library(MASS);library(mvtnorm);library(ggplot2)


#--------------------------------------------------------------------------------#
#                                                                                # 
#  SIMULATE TWO TRAITS WITH CROSS-COVARIANCE MATRICES AND FIT MODELS (figure 2)  #
#                                                                                #
#--------------------------------------------------------------------------------#


# SIMULATE DATA

# simulate tree
n = 200 # number of species
tree <- geiger::sim.bdtree(b=1, d=0, stop="taxa", n=n, extinct=FALSE)
tree <- multi2di(tree) # ensure tree is fully bifurcating

# create phylogenetic covariance matrix, C, from tree.
# C is the expected VCV matrix of a trait for n x n species assuming Brownian motion, 
# corr = T means the correlation matrix rather than covariance matrix is returned. 
C = ape::vcv.phylo(tree, corr = T) 

# identity matrix
I = diag(n)

# number of traits
k = 2

# fixed effects (intercepts for each trait)
u = c(0,0)

# B is the phylogenetic trait-level covariance matrix,
# i.e., B specifies the phylogenetic variance in each trait (diagonals) as well as the phylogenetic covariance between traits (off-diagonals).
# B is k x k (2 x 2 for 2 traits), and specified here as having off-diagonal elements of cov.b = 0.7. (different values supplied to produce the different simulation conditions)
# To construct B, we first define its (co)variance components.
sig.b <- c(b11 = 1, b22 = 1) # sqrt of the phylogenetic variances for each trait; set to 1 so sigma = sigma^2
cor.b = 0.7 # phylogenetic correlation coefficient between traits 1 and 2
Bcor <- matrix(c(c(1,cor.b), c(cor.b,1)),k,k, byrow = T) # phylogenetic correlation matrix; 2 x 2 with off-diagonals set at 0.7
B <- matrix(kronecker(sig.b, sig.b),k,k)*Bcor # Construct B as point-wise product. 
# N.B. Kronecker used in line above just for ease of matrix formatting. Do not confuse with Kronecker
# operation used for generating the cross-correlation matrices for the random effects and residuals below

# E is the residual trait-level covariance matrix
sig.e <- c(e11 = 1, e22 = 1) # sqrt of the residual variance for each trait
cor.e = 0.7 # residual correlation coefficient
Ecor <- matrix(c(c(1,cor.e),c(cor.e,1)),k,k, byrow = T)  # residual correlation matrix; off-diagonals set at 0.7
E <- matrix(kronecker(sig.e, sig.e),k,k)*Ecor # Construct E as point-wise product. 

# simulate phylogenetic random effects and extract for each trait from the vector, b.
b = mvrnorm(n = 1, mu = rep(0,n*k), Sigma = kronecker(B,C))
# n = 1 in order to generate 1 random effect per taxon, mu = rep(0,n*k) centers random effects for all taxa on zero, 
# Sigma = kronecker(B,C) defines the cross-correlation matrix of the random effects. In the kronecker, phylogenetic trait-level covariance captured by B, taxon-level covariance captured by C.
b1 <- b[1:n] # effects 1:n relate to trait 1
b2 <- b[(1:n) + n] # effects (1:n)+n relate to trait 2

# simulate errors and extract
e = mvrnorm(1,rep(0,n*k),kronecker(E,I))# In the kronecker, (phylogenetically) independent trait-level covariance captured by E, taxon-level independent error captured by I.
e1 <- e[1:n]
e2 <- e[(1:n) + n]

# construct response traits from each linear predictor
# i.e., for each trait, add b's and e's to the fixed effect u's
y1 = u[1] + b1 + e1
y2 = u[2] + b2 + e2

# generate data frame
species <- tree$tip.label
d <- data.frame(species,y1,y2)
rownames(d) <- tree$tip.label
plot(d$y1, d$y2)


# FIT MODELS
MRPMM <- brm(mvbind(y1, y2) ~ (1|a|gr(species, cov = C)),
             data = d, 
             family = gaussian(), 
             data2 = list(C = C),
             cores=1,
             chains=1, iter = 2000, thin = 1)

PGLS <-  phylolm(y1 ~ y2, phy=tree, model = "BM", data=d)
PGLS_lambda <-  phylolm(y1 ~ y2, phy=tree, model = "lambda", data=d)
OLS <-  lm(y1 ~ y2, data=d)

# compare model outputs
MRPMM
summary(PGLS)
summary(PGLS_lambda)
summary(OLS)

# repeat above procedure nsim times for each condition (i.e., re-specifying cor.b and cor.e) to replicate data in figure 2


#------------------------------------------------------------------------------------------------------------#
#                                                                                                            #
#  SIMULATE BIVARIATE TRAITS FOR TWO CLADES ACCORDING TO BROWNIAN MOTION, PRICE MODEL and MR-PMM (figure 1)  #
#                                                                                                            #
#------------------------------------------------------------------------------------------------------------#


# SIM PRICE AND PLOT (Price sim run first because tree generated used in subsequent sims)

# define function for Price simulation
sim_Price <- function (N, Sigma, trait = F, min.dist = 0.1, r.scalar = 2.125) {
  
  # sample uniformly from an ellipse defined by Sigma
  # radius of ellipse defined as diagonal element of Sigma
  Sigma <- Sigma*-1;diag(Sigma) <- 1
  
  # specify starting positions for each clade
  niche.space <- matrix(NA,2,2);niche.space[1,] <- c(-1, -1);niche.space[2,] <- c(1, 1)
  
  # alternatively, randomly sample starting positions (MRCA) for each clade within ellipse
  # niche.space <- uniformly::runif_in_ellipsoid(1, Sigma, Sigma[1,1]*r.scalar)
  
  # add one new niche at a time subject to the distance condition min.dist
  while (nrow(niche.space) < N){
    new.niche <- uniformly::runif_in_ellipsoid(1, Sigma, Sigma[1,1]*r.scalar)
    temp.niche.space <- rbind(niche.space, new.niche)
    temp.niche.dist <- as.matrix(dist(temp.niche.space))
    if (min(temp.niche.dist[upper.tri(temp.niche.dist)]) > min.dist) {
      niche.space <- temp.niche.space
    } else NULL
  }
  
  # calculate distance matrix on final niche matrix
  niche.dist <- as.matrix(dist(niche.space))
  
  
  ## GROW TREE
  
  # for each entry find the tip with a lower index that is closest. 
  # tips added sequentially, start at 3 as initial tree must be a cherry (i.e. a tree with 2 tips)
  parent <- rep(0,N)
  for (k in 3:N){
    temp <- niche.dist[k,1:(k-1)]
    parent[k] <- which.min(temp)
  }
  
  # initiate a cherry tree
  atree <- rtree(2)
  atree$edge.length <- c(0,0)
  atree$tip.label <- c("t1","t2")
  num.tips = 2
  
  while(num.tips < N){
    
    # wait some random amount of time, i.e. grow all tips
    growth <- rep(0, nrow(atree$edge))
    growth[atree$edge[,2]<=num.tips] <-rep(rexp(1), num.tips)
    atree$edge.length <- atree$edge.length + growth
    
    # add a tip (niche filled by speciation from existing tip with most similar niche)
    node <- parent[num.tips + 1] # parent node defined by euclidean distances generated above
    newlabel <- paste("t",as.character(num.tips + 1), sep="")
    atree <- phangorn::add.tips(atree, newlabel, node)
    num.tips = num.tips + 1
    
  }
  
  # grow tips once more to prevent zero length terminal sisters
  growth <- rep(0, nrow(atree$edge))
  growth[atree$edge[,2]<=num.tips] <-rep(rexp(1), num.tips)
  atree$edge.length <- atree$edge.length + growth
  
  res <- list(niche.space, atree)
  if(trait) return(res)
  return(atree)
  
}

N = 60
Sigma = matrix(c(1,0.5,0.5,1),2,2) # Sigma defines the shape and scale of the ellipse (ecological attractor) within the Price sim
d.p <- sim_Price(N, Sigma = Sigma, trait = T, r.scalar = 1.5)

# extract data and tree from sim
tree <- d.p[[2]] # use Price generated tree for all subsequent sims
datPrice <- as.data.frame(d.p[[1]])
datPrice$taxa <- tree$tip.label

# Identify clades arising from t1 and t2
phy <- extract.clade(tree, N+2)$tip.label
# assign each taxon to their respective clades
if ("t1" %in% phy){datPrice$clade <- ifelse(datPrice$taxa %in% phy, "phy2","phy1")} else {datPrice$clade <- ifelse(datPrice$taxa %in% phy, "phy1","phy2")}
names(datPrice) <- c("y1", "y2","taxa", "clade")
table(datPrice$clade) # N.B. not necessarily equal (depends on sequence of sampling within ellipse for a particular sim)

# create data frame defining ellipse to pass to stat_ellipse
n <- 10000
r <- as.data.frame(mvrnorm(n, c(0, 0), matrix(c(1,0.5,0.5,1), nrow = 2)*1));r$cor <- "res"
cor <- data.frame(x=r[,1],y=r[,2])
names(cor)[1:2] <- c('y2','y1')

n <- 30
cols <- c("#E69F00","#56B4E9")
p2 <- ggplot(datPrice, aes(x=y2, y=y1, col=clade)) + 
  stat_ellipse(data=cor, col="black", linetype=2) + 
  theme_classic() + 
  scale_color_manual(values=c(cols[2:1])) +
  xlim(-3,3) + ylim(-3,3) + 
  xlab("B") + ylab("A") +
  theme(legend.position = "none") +
  geom_point(alpha=c(rep(1,2),rep(0.75,2*(n-1))),
             size=c(rep(5,2),rep(3,2*(n-1))),
             shape=20)


## BM SIM AND PLOT

# re-scale Price tree
tree$edge.length <- tree$edge.length/max(nodeHeights(tree))
# split Price tree into major clades
phy1 <- keep.tip(tree, datPrice[datPrice$clade=="phy1",]$taxa)
phy2 <- keep.tip(tree, datPrice[datPrice$clade=="phy2",]$taxa)

# simulate two traits via BM on each clade (specifying ancestral trait values for each clade as before) 
phy1_y1<-fastBM(phy1, a = 1, sig2 = 0.5)
phy1_y2<-fastBM(phy1, a = 1, sig2 = 0.5)
phy2_y1<-fastBM(phy2, a = -1, sig2 = 0.5)
phy2_y2<-fastBM(phy2, a = -1, sig2 = 0.5)

datBM <- data.frame(y1 = c(phy1_y1, phy2_y1), y2 = c(phy1_y2, phy2_y2), clade = c(rep("phy1",length(phy1$tip.label)), rep("phy2",length(phy2$tip.label))))
plot(datBM$y1,datBM$y2)

p1 <- ggplot(datBM, aes(x=y2, y=y1, colour=clade)) +
  geom_point(alpha=0.75,size=3,shape=20) +
  scale_color_manual(values=c(cols[2:1])) +
  xlim(-3,3) + ylim(-3,3) +
  xlab("B") + ylab("A") +
  geom_point(aes(x=-1,y=-1),colour=cols[1], size = 3) +
  geom_point(aes(x=1,y=1),colour=cols[2], size = 3) +
  theme_classic() +
  theme(legend.position = "none")


## MR-PMM SIM AND PLOT

# loop over MR-PMM sim to produce data for two clades
species <- c()
y1 <- c()
y2 <- c()
u.mat = matrix(c(1,-1,1,-1), nrow = 2)
trees <- list(phy1,phy2)
for (i in 1:2){
tree.temp <- trees[[i]]
n = length(tree.temp$tip.label)
C = ape::vcv.phylo(tree.temp, corr = T) 
I = diag(n)
k = 2
u = u.mat[i,]
# phy covariance
sig.b <- c(b11 = 0.5, b22 = 0.5)
cor.b = 0.7
Bcor <- matrix(c(c(1,cor.b), c(cor.b,1)),k,k, byrow = T)
B <- matrix(kronecker(sig.b, sig.b),k,k)*Bcor
# ind covariance
sig.e <- c(e11 = 0.5, e22 = 0.5)
cor.e = 0.7
Ecor <- matrix(c(c(1,cor.e),c(cor.e,1)),k,k, byrow = T) 
E <- matrix(kronecker(sig.e, sig.e),k,k)*Ecor
# phy effects
b = mvrnorm(n = 1, mu = rep(0,n*k), Sigma = kronecker(B,C))
b1 <- b[1:n]
b2 <- b[(1:n) + n]
# ind effects
e = mvrnorm(1,rep(0,n*k),kronecker(E,I))
e1 <- e[1:n]
e2 <- e[(1:n) + n]
# construct response traits
y1. = u[1] + b1 + e1
y2. = u[2] + b2 + e2

# add to vector
y1 <- c(y1, y1.)
y2 <- c(y2, y2.)

}

datMRPMM <- data.frame(clade = c(rep("phy1",length(phy1$tip.label)), rep("phy2",length(phy2$tip.label))),y1,y2)

p3 <- ggplot(datMRPMM, aes(x=y2, y=y1, colour=clade)) +
  geom_point(alpha=0.75,size=3,shape=20) +
  scale_color_manual(values=c(cols[2:1])) +
  xlim(-3,3) + ylim(-3,3) + 
  xlab("B") + ylab("A") +
  geom_point(aes(x=-1,y=-1),colour=cols[1], size = 3) +
  geom_point(aes(x=1,y=1),colour=cols[2], size = 3) +
  theme_classic() +
  theme(legend.position = "none")


# combine plots
cowplot::plot_grid(p1, p2, p3, ncol=3, labels = "AUTO")
