## Supplementary 1 for "Phylogenetically conservative trait correlation: quantification and interpretation"

### Supplementary 1: Simulations

For each set of conditions given in Fig 2, we simulated 400 datasets of bivariate trait data. Code is provided as Supplementary 3. First, we generated 400 random bifurcating pure birth trees of 200 species using the sim.bdtree function in the r package 'geiger'. A scaled covariance matrix for each tree was computed using the vcvc.phylo function in the R package 'ape'. We simulated the traits A and B according to the bivariate PMM

$$\begin{pmatrix} A \\ B \end{pmatrix} = \begin{pmatrix} \mu_A & + & b_A & + & e_A \\ \mu_B & + & b_B & + & e_B \end{pmatrix}$$

where $\mu_A = \mu_B = 0$ represent intercept values for each trait. We simulated phylogenetic (b) and residual (e) effects drawn from bivariate normal distributions:

$$(b_A, b_B) \sim N(0, \Sigma^{phy} \otimes C)$$
$$(e_A, e_B) \sim N\left(0, \Sigma^{ind} \otimes I\right)$$

where C is the scaled covariance matrix of the tree, I is the identity matrix, and $\otimes$ is the Kronecker product. We then fit MV-PMM and three GLS models (PGLS with lambda =1, PGLS with lambda fitted and OLS) to each data/tree combination. Relevant parameter estimates from each model fit are presented in figure 2.

The panels illustrating different example outcomes for different models of evolution in Fig 1 were produced by three separate simulations. For panel B, we simulated data following the model outlined in Price (1997). Briefly, an ellipse matching the 95% confidence region of the joint residual density for traits (A, B) assuming a correlation of 0.5 was drawn in trait space to circumscribe possible (i.e. ecologically competent) combinations of the traits A and B. The evolution of two clades was simulated by first defining a cherry tree and two starting positions in trait space (i.e., ancestral states) of (-1, -1) and (1, 1) for the orange and blue clades, respectively. New species were then added (and the tree grown) one species at a time, by uniformly sampling within the ellipse and attaching new species as bifurcations to the nearest taxon in trait space, thereby simulating trait evolution by phylogenetic niche conservatism. A minimum distance in trait space between existing and candidate species was also enforced *sensu* Price (1997). The clades from the resulting tree were used to derive the C matrices for the simulations in panel A and C. For panel A, we simulated two traits via Brownian motion BM for each of the two clades, using the same ancestral values of traits A and B for each clade. This is equivalent to simulating BM on a tree representing two clades appended to long basal branches to create a deep split between the clades. For illustrative purposes, we chose the former approach of simulating data for each clade separately to force the most recent common ancestor MRCA for orange and blue clades to be the same for each panel A-C. For panel C, using the same trees, we simulated values for A and B jointly for each clade from the full MR-PMM structure, (outlined above, also see box 3), setting $\Sigma_{AA}^{phy} = \Sigma_{BB}^{phy} = 1,\ \Sigma_{AA}^{ind} = \Sigma_{BB}^{ind} = 1,\ \rho_{AB}^{ind} = \rho_{AB}^{phy} = 0.7$.
