## Supplementary 2 for "Phylogenetically conservative trait correlation: quantification and interpretation"

**Manuscript title**: Phylogenetically conservative trait correlation: quantification and interpretation

### Mathematical aspects of phylogenetic mixed models

Here we examine some mathematical aspects of phylogenetic mixed models related to the issue of phylogenetic confounding: when a predictor variable contains phylogenetic signal and is thereby correlated with the phylogenetic random effects (or equivalently, the phylogenetic component of total residual variance). Phylogenetic confounding typically occurs in single-response models such as PGLS when traits are included as predictors.

We study the general form of a Gaussian multi-response phylogenetic mixed model which includes parameters to estimate the phylogenetic signal in the variances of each response trait as well as the covariances of each trait pair. We specialise to the case of two responses (i.e., two traits A and B) with the aim to investigate how terms in the multi-response model manifest in the conditional model $A|B$. Expressed in its marginal form—having integrated out the phylogenetic random effects—the multi-response phylogenetic mixed model is

$$\begin{pmatrix} A \\ B \end{pmatrix} \sim \mathrm{N}\left( \begin{pmatrix} \mu_A \\ \mu_B \end{pmatrix}, \Sigma^{\mathrm{phy}} \otimes C + \Sigma^{\mathrm{ind}} \otimes I \right), \tag{1}$$

where $\mu_A$ and $\mu_B$ are fixed effects, $C$ is the phylogenetic relatedness matrix, $I$ is the identity matrix, and

$$\Sigma_{ij}^{\mathrm{phy}} = \sigma_i^{\mathrm{phy}} \sigma_j^{\mathrm{phy}} \rho_{ij}^{\mathrm{phy}} \quad \text{and} \quad \Sigma_{ij}^{\mathrm{ind}} = \sigma_i^{\mathrm{ind}} \sigma_j^{\mathrm{ind}} \rho_{ij}^{\mathrm{ind}}, \tag{2}$$

for $i, j = A, B$ are trait-level covariances. The vectors $A$, $B$, $\mu_A$ and $\mu_B$ have length $n$ and the matrices $C$ and $I$ are of dimension $n \times n$ where $n$ is the number of taxa (see Box 3 in the main manuscript for further details).

From the theory of multivariate normal distributions, the conditional distribution of $A$ given $B$ is

$$A|B \sim \mathrm{N}\left(\mu_{A|B}, \Sigma_{A|B}\right) \tag{3}$$

where

$$\mu_{A|B} = \mu_A + \Sigma_{AB}\Sigma_{BB}^{-1}(B - \mu_B) \tag{4}$$

$$\Sigma_{A|B} = \Sigma_{AA} - \Sigma_{AB}\Sigma_{BB}^{-1}\Sigma_{BA} \tag{5}$$

for

$$\Sigma_{AB} = \sigma_A^{\mathrm{phy}} \sigma_B^{\mathrm{phy}} \rho_{AB}^{\mathrm{phy}} C + \sigma_A^{\mathrm{ind}} \sigma_B^{\mathrm{ind}} \rho_{AB}^{\mathrm{ind}} I \qquad (= \Sigma_{BA}) \tag{6}$$

$$\Sigma_{BB} = (\sigma_B^{\mathrm{phy}})^2 C + (\sigma_B^{\mathrm{ind}})^2 I. \tag{7}$$

The conditional mean $\mu_{A|B}$ is the best (in the sense of least squared error for one response and maximal correlation for multiple responses) linear predictor of $A$ as a function of $B$. The presence of phylogenetic signal in both $A$ and $B$ means that the regression matrix

$$\beta_{A|B} := \Sigma_{AB}\Sigma_{BB}^{-1}. \tag{8}$$

generally comprises non-zero off-diagonal terms rather than being a scalar multiple of the identity matrix as it is constrained to be in standard single-response models such as OLS and PGLS.

To investigate phylogenetic confounding and to facilitate comparison of the conditional model (3) with the parameter estimates of single-response models, we derive a series representation of $\beta_{A|B}$. Viewed as a function of $C$, $\Sigma_{BB}^{-1}$ can be expressed as a matrix series expansion about the identity:

$$\begin{aligned}\Sigma_{BB}^{-1}(C) &= \left((\sigma_B^{\text{phy}})^2 C + (\sigma_B^{\text{ind}})^2 I\right)^{-1} \\ &= \sum_{n=0}^{\infty} c_n (C-I)^n \end{aligned} \tag{9}$$

for

$$\begin{aligned} c_n &= \frac{1}{n!}\,\frac{\mathrm{d}^n}{\mathrm{d}C^n}\Sigma_{BB}^{-1}(C)\bigg|_{C=I} \\ &= \frac{(-1)^n\left((\sigma_B^{\text{phy}})^2\right)^n}{\left((\sigma_B^{\text{phy}})^2 + (\sigma_B^{\text{ind}})^2\right)^{n+1}} \\ &= \frac{(-\lambda_B)^n}{\sigma_B^2}, \end{aligned} \tag{10}$$

where

$$\lambda_B = \frac{(\sigma_B^{\text{phy}})^2}{(\sigma_B^{\text{phy}})^2 + (\sigma_B^{\text{ind}})^2} \tag{11}$$

and

$$\sigma_B^2 = (\sigma_B^{\text{phy}})^2 + (\sigma_B^{\text{ind}})^2. \tag{12}$$

The leading terms in the series (9) are:

$$\Sigma_{BB}^{-1}(C) = \frac{1}{\sigma_B^2} I - \frac{\lambda_B}{\sigma_B^2}(C-I) + \frac{\lambda_B^2}{\sigma_B^2}(C-I)^2 - \cdots \tag{13}$$

which converges provided $\lambda_B|C-I| < 1$. Writing

$$\Sigma_{AB} = (\sigma_A^{\text{phy}}\sigma_B^{\text{phy}}\rho_{AB}^{\text{phy}} + \sigma_A^{\text{ind}}\sigma_B^{\text{ind}}\rho_{AB}^{\text{ind}})(\lambda_{AB}C + (1-\lambda_{AB})I) \tag{14}$$

for

$$\lambda_{AB} := \frac{\sigma_A^{\text{phy}} \sigma_B^{\text{phy}} \rho_{AB}^{\text{phy}}}{\sigma_A^{\text{phy}} \sigma_B^{\text{phy}} \rho_{AB}^{\text{phy}} + \sigma_A^{\text{ind}} \sigma_B^{\text{ind}} \rho_{AB}^{\text{ind}}}, \tag{15}$$

we substitute (13) and (14) into (8) to obtain

$$\beta_{A|B} = \frac{\sigma_A^{\text{phy}} \sigma_B^{\text{phy}} \rho_{AB}^{\text{phy}} + \sigma_A^{\text{ind}} \sigma_B^{\text{ind}} \rho_{AB}^{\text{ind}}}{\sigma_B^2} \left( I + (\lambda_{AB} - \lambda_B) \sum_{i=1}^{\infty} (-\lambda_B)^{i-1} (C - I)^i \right). \tag{16}$$

Let us also consider a decomposition of the 'variance in $A$ explained by $B$' which is the difference $\Sigma_{AA} - \Sigma_{A|B}$. Using (5) and (14), we write this as

$$\Sigma_{AB} \Sigma_{BB}^{-1} \Sigma_{BA} = \frac{(\sigma_A^{\text{phy}} \sigma_B^{\text{phy}} \rho_{AB}^{\text{phy}} + \sigma_A^{\text{ind}} \sigma_B^{\text{ind}} \rho_{AB}^{\text{ind}})^2}{\sigma_B^2} \Sigma_{\lambda_{AB}} \Sigma_{\lambda_B}^{-1} \Sigma_{\lambda_{AB}} \tag{17}$$

where

$$\Sigma_{\lambda_{AB}} = \lambda_{AB} C + (1 - \lambda_{AB}) I \quad \text{and} \quad \Sigma_{\lambda_B} = \lambda_B C + (1 - \lambda_B) I. \tag{18}$$

For the simple case $\sigma_A^{\text{phy}} = \sigma_B^{\text{phy}} = 0$, the conditional variance is $\Sigma_{A|B} = \Sigma_{AA}(1 - (\rho_{AB}^{\text{ind}})^2)$; similarly for the case $\sigma_A^{\text{ind}} = \sigma_B^{\text{ind}} = 0$.

The issue of phylogenetic confounding for conditional regression models $A|B$ arises due to the additive decomposition of the total variance in (1) and in particular the partitioned component (6). The addition of the independent and phylogenetic covariances expressed as $(\sigma_A^{\text{phy}} \sigma_B^{\text{phy}} \rho_{AB}^{\text{phy}} + \sigma_A^{\text{ind}} \sigma_B^{\text{ind}} \rho_{AB}^{\text{ind}})$ factors both the regression matrix $\beta_{A|B}$ (16) and the variance explained $\Sigma_{AB} \Sigma_{BB}^{-1} \Sigma_{AB}$ (17) showing that regression of $A$ on $B$ confounds the slope estimate with the phylogenetically structured residual variance. When the independent and phylogenetic covariance components differ in sign, they will have a negating effect on the magnitude of the slope masking these distinct albeit competing forces on trait coevolution. For strictly predictive goals, this confounding is likely inconsequential, however, for inferential goals it is problematic because the attribution of variance explained in $A$ amongst predictor and phylogeny is not identifiable. This confounding impacts on the interpretation and validity of hypothesis tests based on the slope.

The series expansion (16) shows that the leading scalar term is 'corrected' by powers of $(C - I)$ as a function of $\lambda_{AB} - \lambda_B$: the difference between the phylogenetic signal in the variance of $B$ and that in the covariance of $A$ and $B$. In other words, single-response models with 'scalar' regression matrices such as PGLS can only adequately approximate the full conditional multi-response model when this difference in signal is small. We note the simple special cases $\sigma_B^{\text{phy}} = 0$ (ordinary PGLS) and $\sigma_B^{\text{ind}} = 0$ for which only the leading term of (16) is non-zero yielding the more familiar regression coefficients:

$$\beta_{A|B} = \begin{cases} \dfrac{\sigma_A^{\text{phy}} \rho_{AB}^{\text{phy}}}{\sigma_B^{\text{phy}}} I\,; & \text{when } \sigma_B^{\text{ind}} = 0 \ (\lambda_B = 1) \\[2ex] \dfrac{\sigma_A^{\text{ind}} \rho_{AB}^{\text{ind}}}{\sigma_B^{\text{ind}}} I\,; & \text{when } \sigma_B^{\text{phy}} = 0 \ (\lambda_B = 0). \end{cases} \tag{19}$$

For data simulated under the full two-response model, it is not straightforward to compare the slope estimates of single-response models such as OLS and PGLS to the regression matrix $\beta_{A|B}$ of the conditional model due to the former being scalar approximations using misspecified models. A comparison of the coefficient of leading term in the series expansion (16) to the estimated slopes for the four simulations presented in Figure 2 in the manuscript suggests that OLS ($\lambda = 0$) approximates the leading coefficient, PGLS/PIC ($\lambda = 1$) approximates the independent component of the covariance, and that PGLS ($\lambda$ estimated) attains an intermediate value. All of the simulations use the fixed values $\sigma_A^{\text{ind}} = \sigma_B^{\text{ind}} = \sigma_A^{\text{phy}} = \sigma_B^{\text{phy}} = 1$ but differ according to the values of $\rho_{AB}^{\text{ind}}$ and $\rho_{AB}^{\text{phy}}$.
